# Microbial Communities Associated with Sustained Anaerobic Reductive Dechlorination of α-, β-, γ-, and δ-Hexachlorocyclohexane Isomers to Monochlorobenzene and Benzene

**DOI:** 10.1101/770354

**Authors:** Wenjing Qiao, Luz A. Puentes Jácome, Xianjin Tang, Line Lomheim, Minqing Ivy Yang, Sarra Gaspard, Ingrid Regina Avanzi, Jichun Wu, Shujun Ye, Elizabeth A. Edwards

## Abstract

Intensive historical and worldwide use of the persistent pesticide technical-grade hexachlorocyclohexane (HCH), composed of the active ingredient γ-HCH (called lindane) along with several other HCH isomers, has led to widespread contamination. We derived four anaerobic enrichment cultures from HCH-contaminated soil capable of sustainably dechlorinating each of α-, β-, γ-, and δ-HCH isomers stoichiometrically and completely to benzene and monochlorobenzene (MCB). For each isomer, the dechlorination rates increased progressively from <3 *µ*M/day to ∼12 *µ*M/day over two years. The molar ratio of benzene to MCB produced was a function of the substrate isomer, and ranged from β (0.77±0.15), α (0.55±0.09), γ (0.13±0.02) to δ (0.06±0.02) in accordance with pathway predictions based on prevalence of antiperiplanar geometry. Cultivation with a different HCH isomer resulted in distinct bacterial communities, but similar archaeal communities. Data from 16S rRNA gene amplicon sequencing and quantitative PCR revealed significant increases in the absolute abundance of *Pelobacter* and *Dehalobacter*, especially in the α-HCH and δ-HCH cultures. This study provides the first direct comparison of shifts in anaerobic microbial communities induced by the dechlorination of distinct HCH isomers. It also uncovers candidate microorganisms responsible for the dechlorination of α-, β-, γ-, and δ-HCH, a key step towards better understanding and monitoring of natural attenuation processes and improving bioremediation technologies for HCH-contaminated sites.

## 1. Introduction

Hexachlorocyclohexane (HCH) was first synthesized by Michael Faraday in 1825^1^ and has been heavily used as a wide spectrum insecticide since the 1940s.^2^ γ-HCH, also known as lindane, is the only isomer with pesticidal activity. However, technical-grade HCH used as a commercial pesticide consisted of a mixture of α (60%–70%), β (5%–12%), γ (10%–15%), and δ (6%–10%) isomers.^1, 3^ During manufacturing and application of technical-grade HCH, large quantities were improperly disposed,^4^ leading to widespread heavy contamination of soils with all isomers.^5-7^ Although the use of HCH is restricted or completely banned in most countries, HCH isomers are still detected in water, soil, sediments, plants and animals all over the world.^8^ HCH isomers are recalcitrant, bio-accumulating, neurotoxic, teratogenic and carcinogenic substances;^9, 10^ and were included in the Stockholm Convention list of persistent organic pollutants in 2009.^2, 11, 12^

HCH isomers differ in the orientation of the chlorine atom substituents on the cyclohexane ring (Figure S1 in Supplementary Information (SI)), which can either be axial (vertically above or below the ring) or equatorial with the cyclohexane ring. These structural differences give rise to considerably different physical and chemical properties (Table S1), owing to differences in dipole moment and steric effects that influence reactivity, especially for nucleophilic and electrophilic reactions.^3, 13, 14^ Bulky axial substituents are closer to each other on the same side of the ring, a phenomenon known as 1,3-diaxial interaction^15^ which increases the overall energy of the molecule and provides specific sites for enzyme attack.^16, 17^ For example, α-and γ-HCH, having two and three axial chlorine atoms, are more easily biodegraded than δ-HCH containing only one axial chlorine atom.^5, 9, 18^ β-HCH is the least reactive and most persistent isomer because all the chlorine atoms occupy more commodious equatorial positions with lower energy.^9, 18, 19^ In addition, HCH has two different chair conformers which are at rapid equilibrium. The conformers with fewer axial chlorine atoms and lower torsional strain are always favored (Figure S1 and Table S2).

Both abiotic and biotic transformation reactions have been observed with HCH isomers. Abiotic reactions occur under alkaline (pH≥8.3) and chemically- or electrochemically-reducing conditions.^2, 20-24^ Dehydrochlorination and chemical dichloroelimination are the principal abiotic reactions. Dehydrochlorination (i.e., elimination of HCl) from two adjacent carbon atoms (when substituents are in the anti-axial position) does not involve electron transfer and may occur abiotically and spontaneously, while dichloroelimination removes two chlorines from two adjacent carbons with a net input of two electrons to produce an olefin. Dehydrochlorination of HCH isomers has only been observed in alcoholic solvents under alkaline conditions (pH≥8.3), and the major end product was 1,2,4-trichlorobenzene (TCB) with lesser amounts of 1,2,3- and 1,3,5-TCB after three successive steps of dehydrochlorination.^20, 21, 25^ The half-life of γ-HCH decreased from 1100 days at pH 6.9 to 126 days at pH 8.3 and to about 30 min at pH 12.^21^ In contrast, chemical dichloroelimination of γ-HCH is relatively pH insensitive and can be mediated by iron sulfides (FeS) and activated carbon to generate dichlorobenzene (DCB) and TCB isomers as end products.^2, 21, 22^ These reactions were reported to only occur at the surface of FeS and electrons were provided by the solid rather than dissolved in solution phase.^21^ The electrochemical reduction of HCHs to benzene has also been described in detail.^23, 24^ Although the aforementioned abiotic reactions have been demonstrated in the laboratory, in the environment these type of reactions are expected to happen very slowly because pH is generally around neutral and reducing equivalents are not in excess. In fact, the amount of detectable HCH that remains in HCH-contaminated soils varies depending on the characteristics of the contaminated site and the composition of the isomer mixture.^26^ The environmental half-life of β-HCH, for example, is estimated to be in the order of years in contrast to γ-HCH which is in the order of days. Soil moisture, temperature, and the oxidation-reduction potential of the subsurface will also impact the rates and extent of abiotic and biotic transformations in contaminated environments.

HCH isomers can be biodegraded under both aerobic and anaerobic conditions. *Sphingomonas, Sphinobium* and *Pseudomonas* are the dominant genera responsible for aerobic degradation of HCH and use HCH as a sole carbon source.^6, 27-29^ The *lin* genes involved in aerobic HCH isomers degradation pathway have been fully characterized.^6, 10, 30-33^ HCH contamination is also reported for anaerobic environments such as river sediments near former agricultural land and landfills.^34, 35^ Anaerobically, reductive dechlorination of HCH isomers has been observed in soil, sludge, and enrichment cultures.^5, 18, 36-38^ *Clostridium, Citrobacter, Desulfovibrio*, and *Desulfococcus* have been found to mediate the dechlorination of HCH isomers co-metabolically.^10, 16, 39-42^ In these studies, relative dechlorination rates depended on the isomer and were found to be in the order γ > α > δ > β.^10^ An enrichment culture capable of using γ-HCH as a terminal electron acceptor was reported in 2011, yet no known organohalide-respiring genera were detected.^43^ To date, only organohalide-respiring *Dehalobacter* sp. E1 has been reported to dechlorinate β-HCH coupled to growth.^19, 44^ *Dehalococcoides mccartyi* strain 195 was recently reported to dechlorinate γ-HCH and use γ-HCH as a substrate for growth;^45^ however, the specific genes and enzymes responsible for reductive dechlorination of HCH isomers have not been identified. Tetrachlorocyclohexene (TeCCH) isomers have been detected as metabolites.^5, 40, 46, 47^ The dechlorination pathway of β-HCH was proposed to comprise two sequential dichloroelimination reactions followed by either another dichloroelimination to produce benzene or a dehydrochlorination to generate monochlorobenzene (MCB).^5, 19^ These dechlorination processes and the end-products have not been systematically characterized for all HCH isomers.

In this study, a γ-HCH-dechlorinating enrichment culture (GT1) derived from HCH-contaminated soil from Guadeloupe (French West Indies) was used to inoculate a series of transfers into mineral medium that were amended with either α-, β-, γ-, or δ-HCH as the electron acceptor and an acetone/ethanol mix as electron donor. The objectives of this study were: 1) to measure the dechlorination rates and products of α-, β-, γ-, and δ-HCH; 2) to relate the observed dechlorination to the stereochemistry of each HCH isomer; 3) to compare and differentiate the resulting enriched microbial communities; and 4) to identify dechlorinating microorganisms.

## 2. Materials and Methods

### Reagents

Neat α-, β-, δ-and γ-HCH isomers, MCB, benzene, acetone and ethanol were purchased from Sigma-Aldrich at the highest purity available (analytical standard grade for HCH isomers, ≥99% purities for all other chemicals).

### Dechlorinating Enrichment Cultures

In 2010, a series of anaerobic microcosms were set up with soil, sediments and waste from various agricultural locations and streams from the island of Guadeloupe (French West Indies). A largely sediment-free transfer from these original microcosms, called GT1, was enriched and maintained over a period of about 2 years with γ-HCH as electron acceptor. Culture GT1 was used to inoculate (1% by volume) a series of 150-ml bottles containing 90 mL of sterile defined ferrous sulfide-reduced mineral medium.^48^ These cultures were amended with one of the α-, β-, γ- or δ-HCH isomers (designated as α-, β-, γ- and δ-HCH enrichment cultures, Figure 1) and re-amended repeatedly as HCH became depleted. The original microcosms and GT1 culture were fed much less frequently. Sterile controls (with autoclaved culture) were prepared at the same time. All cultures were prepared in triplicates and incubated statically in dark at room temperature in a Coy anaerobic chamber (Coy Laboratory Products, Madison, WI) supplied with a CO_2_/H_2_/N_2_ (10/10/80 percent by volume) gas mix. HCH isomers were initially dissolved in acetone and ethanol to make feeding stock solutions in which the acetone/ethanol mix served as electron donors. Because each isomer has a different solubility in this solvent mix, the feeding stocks had different concentrations of HCH, and this led to adding different amounts of electron donors to each set of enrichments. Specifically, donor was added at very high excess to the β-HCH and α-HCH bottles: 511 and 201 times the electron equivalents (eeq) required for complete dechlorination, respectively. Donor was added at lower dosages for the δ- and γ-HCH bottles, i.e., 98 and 46 times eeq, respectively. To harmonize and reduce donor addition, a solvent-evaporation method was devised to add HCH isomer to each culture bottle (without solvent) and this method was implemented from Day 499 onwards (see “HCH and electron donor amendment methods” in the SI). With the new method, only ethanol was provided as donor at a consistent 10 times the eeq required for complete dechlorination. Dechlorination rates were calculated assuming zero-order kinetics as 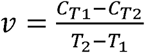, where *v* is the dechlorination rate (*µ*M/day) and *C*_*T1*_ and *C*_*T2*_ are the molar concentrations at times *T*_*1*_ and *T*_*2*_, respectively. The recovery of products based on the amount of HCH isomer amended was calculated as 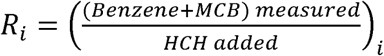, where compound names represent molar concentrations and *i* represents the specific isomer: α-, β-, γ- or δ-HCH. The cultures were re-fed when the MCB and benzene produced were approximately equal to the amount of HCH amended. The amounts of HCH isomers and electron donor amended to each culture are listed in Table S3.

**Figure 1.**
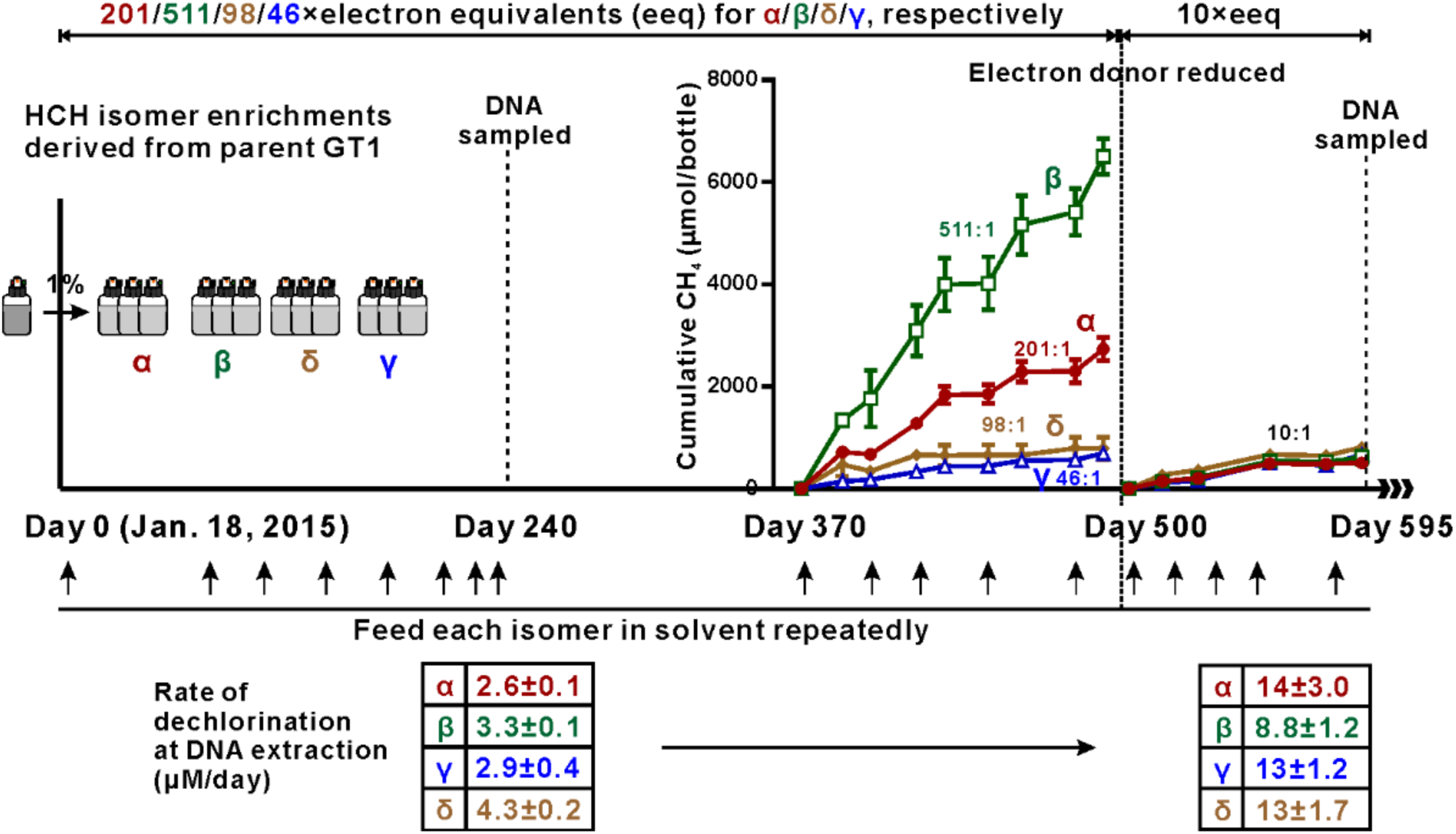
Event timeline for HCH dechlorinating enrichment cultures. Notes: The original inoculum was a γ-HCH-amended enrichment called GT1, derived from Guadeloupe soil microcosms set up in 2010. The γ-HCH dechlorinating triplicate culture set was set up 130 days later than the other three culture sets. DNA samples were collected on Day 240 and 595. On Day 499, the feeding method was changed so that the same amount of donor was supplied to all isomer bottles, reducing the amount of donor added. Corresponding electron equivalent ratios of donor to HCH are shown for each isomer; the ratio was harmonized to 10:1 after day 499. The short arrows show feeding events (see Table S3 for detailed feeding information).

### Analytical Methods

MCB and benzene concentrations were quantified by an Agilent 7890 GC-FID equipped with an Agilent J&W GS-Q column (30 m × 0.53 mm). Aqueous culture samples (1 mL) were added to 5 mL of acidified deionised water (∼pH = 2) and then equilibrated in an Agilent G1888 autosampler at 70 °C for 40 min. After equilibration, 3 mL of headspace sample was injected into GC via a packed inlet. The temperature of injector and detector were 200 °C and 250 °C, respectively. The oven temperature program was initially set at 35 □ for 1.5 min, increased to 100 °C at 15 °C/min, then increased to 185 °C at 5 °C/min, then increased to 200 °C at 20 °C/min and finally held at 200 °C for 10 min.

### DNA Extraction and Quantitative PCR (qPCR)

Well mixed samples (10 mL) from parent culture (GT1), and the α-, β-, γ- and δ-HCH enrichment cultures were collected after 240 and 595 days of incubation and spun down at 10,000xg for 15 min at room temperature. The pellets were used for DNA extraction using the DNeasy PowerSoil kit (QIAGEN) according to the manufacturer’s protocol except that DNA was eluted in 50 *µ*L of sterile UltraPure distilled water (Invitrogen Carlsbad, CA). A NanoDrop ND-1000 spectrophotometer (NanoDrop Technologies, Wilmington, DE, USA) was used to assess DNA concentrations and quality. Total bacteria and archaea were enumerated using published qPCR primers sets.^49, 50^ Serial dilutions of plasmids containing corresponding 16S rRNA gene fragments were used to generate standard curves (see “Quantitative PCR” in SI for additional details). Each reaction (20 *µ*L) was run in duplicate. All the reactions were prepared in a PCR UV cabinet (ESCO Technologies, Hatboro, PA). The analyses were performed with a Bio-Rad C1000 thermocycler (Bio-Rad, Hercules, California, USA) and the CFX Manager software.

### 16S rRNA Gene Amplicon Sequencing and Data Analysis

DNA samples were sequenced at McGill University and Genome Quebec Innovation Center. The original DNA extracted from the parent GT1 culture was sequenced using the Roche GS FLX Titanium platform with the primers 926f (5’-AAACTYAAAKGAATTGACGG-3’) and 1392r (5’-ACGGGCGGTGTGTRC-3’),^51, 52^ while DNA extracted from the enrichment cultures on Day 240 and 595 were sequenced using the Illumina MiSeq PE300 platform with slightly modified primers, i.e., 926f modified (5’-AAACTYAAAKGAAT**W**G**R**CGG-3’), and 1392r modified (5’-ACGGGCGGTG**W**GTRC-3’), which were derived from previous publications.^51, 53^ Please refer to “Amplicon Sequencing” in SI for additional details.

The raw sequence data was processed using the Quantitative Insights Into Microbial Ecology (QIIME v1.9.1) pipeline^54^ with default settings unless stated otherwise. The sequences obtained from the Roche GS FLX Titanium platform with read length > 250 bp, quality score > 27 and homopolymer length < 8 bp, were demultiplexed and analyzed. The forward and reverse raw reads obtained from Illumina sequencing were joined with 50 bp overlap using the default mismatch (8%). The joined reads were then quality filtered (quality score >19). All the reads from Pyrotag and Illumina sequencing were merged for downstream analysis. Chimera checking was performed using the Usearch 61^55^ algorithm and the Ribosomal Database Project (v11) as reference.^56, 57^ The non-chimeric sequences were further clustered into distinct 16S rRNA gene-based Operational Taxonomic Units (OTUs) based on 97% similarity using usearch 61^55^ and the Greengenes Database (v13.8).^58^ The most abundant sequence within each cluster was picked as the representative sequence for each OTU.

### Ordination Analysis

To compare microbial communities among the cultures receiving different HCH isomers, ordination analyses were conducted. Because of excess acetone and/or ethanol provided as donor, archaea flourished in the microbial community. Therefore, in an attempt to better delineate differences between the bacterial or archaeal communities, archaea and bacteria data were separated for the ordination analysis. Nonmetric multidimensional scaling (NMDS) was performed using the metaMDS function, Vegan Package v2.5-5 in R (v3.6.1) with absolute abundance of bacteria or archaea calculated by multiplying total bacteria or archaea gene copies (obtained from qPCR) by the normalized relative abundance within bacterial or archaeal OTUs obtained from amplicon sequencing. The Bray-Curtis dissimilarity index was used to calculate bacterial or archaeal community phylogenetic distance, which was then visualized using the NMDS numerical technique. The number of dimensions was chosen to minimize stress which is a measure of the agreement between dissimilarity scores and predicted ordination distances.^59, 60^

## 3. Results and Discussion

### 3.1 Reductive Dechlorination of HCH Isomers

Dechlorination of all four HCH isomers was sustained over 600 days. The average recovery, *R*_*i*_, of the four isomers in this study in the active enrichments was 92%±5% (N=12), confirming that each HCH isomer was stoichiometrically and completely transformed to MCB and benzene. To view comprehensive time course data, the cumulative production (rather than concentration) of the end-products MCB and benzene was plotted over time (Figure 2), providing an unprecedented recording of long-term dechlorination profiles for each of the HCH isomers. Over time, the rates of dechlorination in the α-, β-, γ- and δ-HCH cultures increased from 2.6 ± 0.11, 3.3 ± 0.13, 2.9 ± 0.42, and 4.3 ±0.18 *µ*M/day on Day 240, to 14 ± 3, 8.8 ± 1.2, 13 ± 1.2 and 13 ± 1.7 *µ*M/day on Day 595, respectively. In all cases, dechlorination rates increased progressively, strongly suggesting that the dechlorination processes were growth-linked. In sterile controls, abiotic dechlorination of HCH to benzene and MCB was insignificant relative to biotic rates (Figure S2).

**Figure 2.**
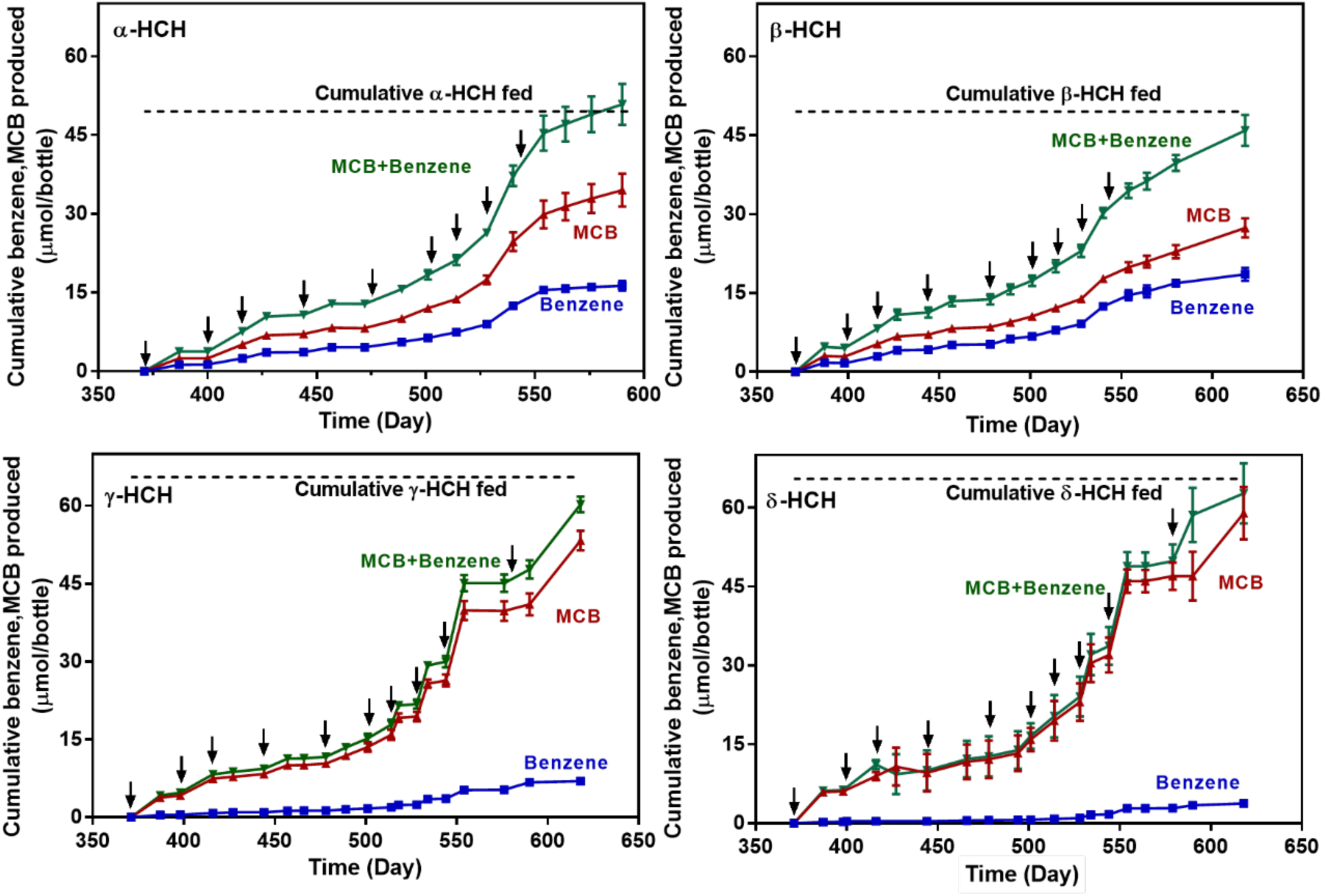
Cumulative dechlorination of α-, β-, γ- and δ-HCH to MCB and benzene in triplicate enrichment cultures over a 250-day window (Days 370-620). Donor amendment was harmonized to 10 times electron equivalents from Day 500 onwards and the amount of HCH fed was increased gradually. The data shown corresponds to the mean of triplicate bottles and the error bars correspond to one +/- standard deviation of the mean. The dashed line represents the total amount of HCH added to each bottle in a set. HCH was re-amended only when the sum of MCB and benzene equaled the amount added. Black arrows represent feeding events (see Table S3 for detailed feeding information). Data from Day 370 to Day 620 are shown; prior data was not recorded as frequently. DNA samples were taken on Days 240 and 595.

The ratio of benzene to MCB produced varied by the isomer type and was relatively consistent as follows: β (0.77±0.15) > α (0.55±0.09) > γ (0.13±0.02) > δ (0.06±0.02) (Figure 3). These ratios were comparable to values published previously (comparison shown in Table S4). Among previous reports, only Doesburg et al. (2005) compared more than one isomer and found that the ratios decreased in the same order as what we found, i.e., β > α > γ.^19^ They hypothesized that the ratio decreased with increasing number of axial chlorine atoms.^19^ However, Doesburg et al. did not test the δ-isomer which has only one axial chlorine atom. In this study, we found that the δ isomer has the lowest ratio, contradicting Doesburg’s hypothesis. Rather, we observed that the α- and β-HCH with an even number of equatorial chlorine atoms produced more benzene than γ- and δ-HCH with an odd number of equatorial chlorine atoms, which is in accordance with the predicted reactivity of these isomers (see below).

**Figure 3.**
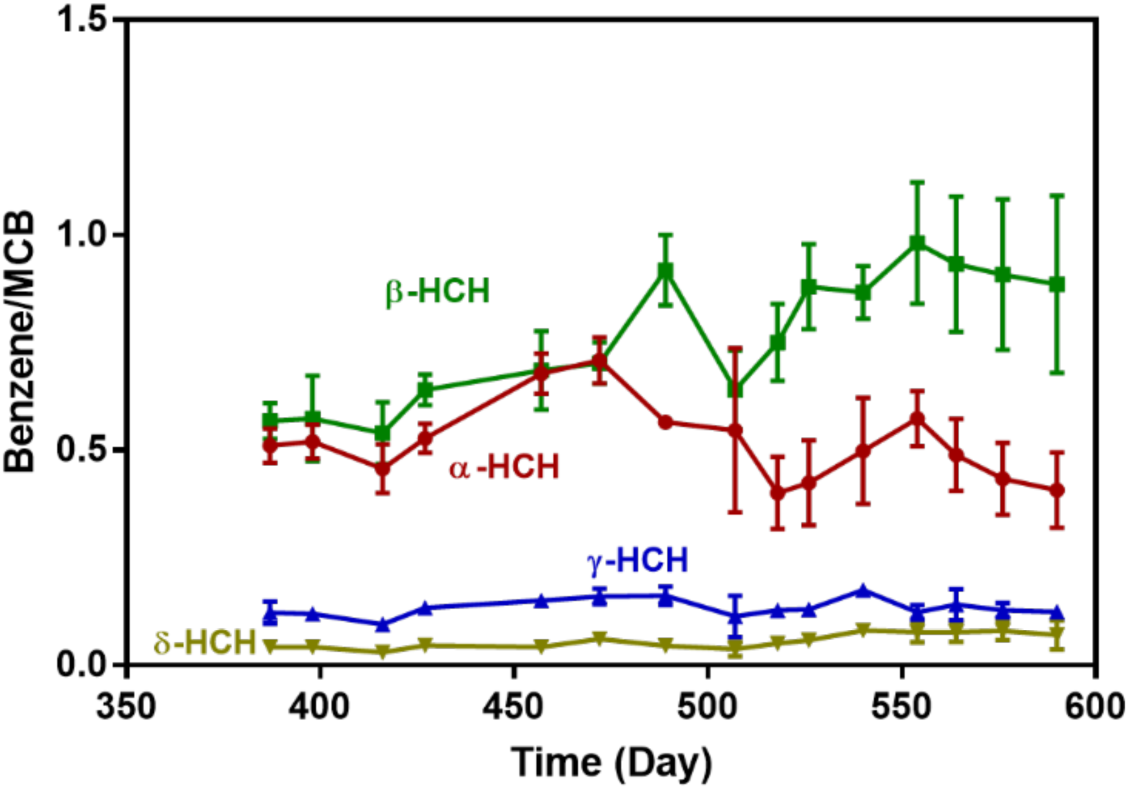
Measured ratios of benzene to MCB in each HCH isomer enrichment culture over time. The data are average of triplicate bottles and the error bars present one +/- standard deviation of the mean.

### 3.2 Dechlorination Pathway Predictions and Benzene to MCB Product Ratios

As introduced previously, abiotic dehydrochlorination reactions (eliminating HCl) can occur under alkaline conditions producing TCB isomers as the major end products.^20, 21, 25^ Moreover, Liu et al. reported that the reducing agent FeS enhanced dichloroelimination of γ-HCH at a concentration of 10 g/L.^21^ In this study, FeS was added as a reducing agent at much lower concentration (< 0.02 g/L), presumably too low to provide sufficient surface area for reaction because we observed that abiotic reactions were insignificant (Figure S2). We also observed stoichiometric production of MCB and benzene, indicating that either two or three biologically-mediated reductive dichloroelimination steps must have occurred. We sketched the most likely sequential elimination reaction pathways for each HCH isomer based on the prevalence of antiperiplanar geometry. Antiperiplanar geometry is where leaving groups (Cl or H atoms) on adjacent carbon atoms are in axial positions such that the all atoms are co-planar, thus facilitating the formation of a new π bond. We compared these predicted pathways with observed benzene to MCB ratios (Figure 4). For all isomers, the first reaction can be a reductive dichloroelimination reaction to produce TeCCH (Figure 4). For the β- and δ-HCH, a ring flip from the most stable conformation must occur first to obtain adjacent axial anti-parallel (or antiperiplanar) chlorines. For β-HCH, in which all chlorine atoms are axial after ring flip alternating on either side of the ring, three successive antiperiplanar reductive dichloroelimination reactions via TeCCH and dichlorocyclohexadiene (DCCH) and to benzene are possible. Accordingly, this isomer had the highest observed benzene yield. Three successive dichloroelimination reactions with proper geometry are also possible with α-HCH (Figure 4), although there are only 4 reactive positions initially, instead of 6 with β-HCH. The α-HCH isomer exists as two enantiomers and thus produces two different TeCCH molecules. In fact, this was reported in a previous study in which two different GC-MS peaks of the intermediate TeCCH were observed during the transformation of α-HCH.^5^ The TeCCH isomers produced from β- and γ-HCH were also detected previously.^5, 40, 46^ In TeCCH, the single bonded carbon atoms are still chiral. TeCCH is not stable as HCH because aromatization is faster with one or two double bonds already in the ring.^46^ TeCCH can undergo either dichloroelimination or spontaneous dehydrochlorination depending on the orientation of substituents. The TeCCH produced by the γ- and δ-HCH isomers would favor dehydrochlorination rather than dichloroelimination, resulting in more MCB production compared to α- and β-HCH. The dichloroelimination of the *trans* vicinal chlorine atoms of TeCCH generated a chemically labile compound, DCCH. The transformation of DCCH to MCB or benzene was previously assumed to be spontaneous, given the strong electromeric effect (i.e. resonance stabilization) associated with the formation of the aromatic ring.^25, 61^ The *trans-*DCCH produced by α- and β-HCH, predictably generates more benzene than *cis-*DCCH owing to favored anti-elimination reactions.

**Figure 4.**
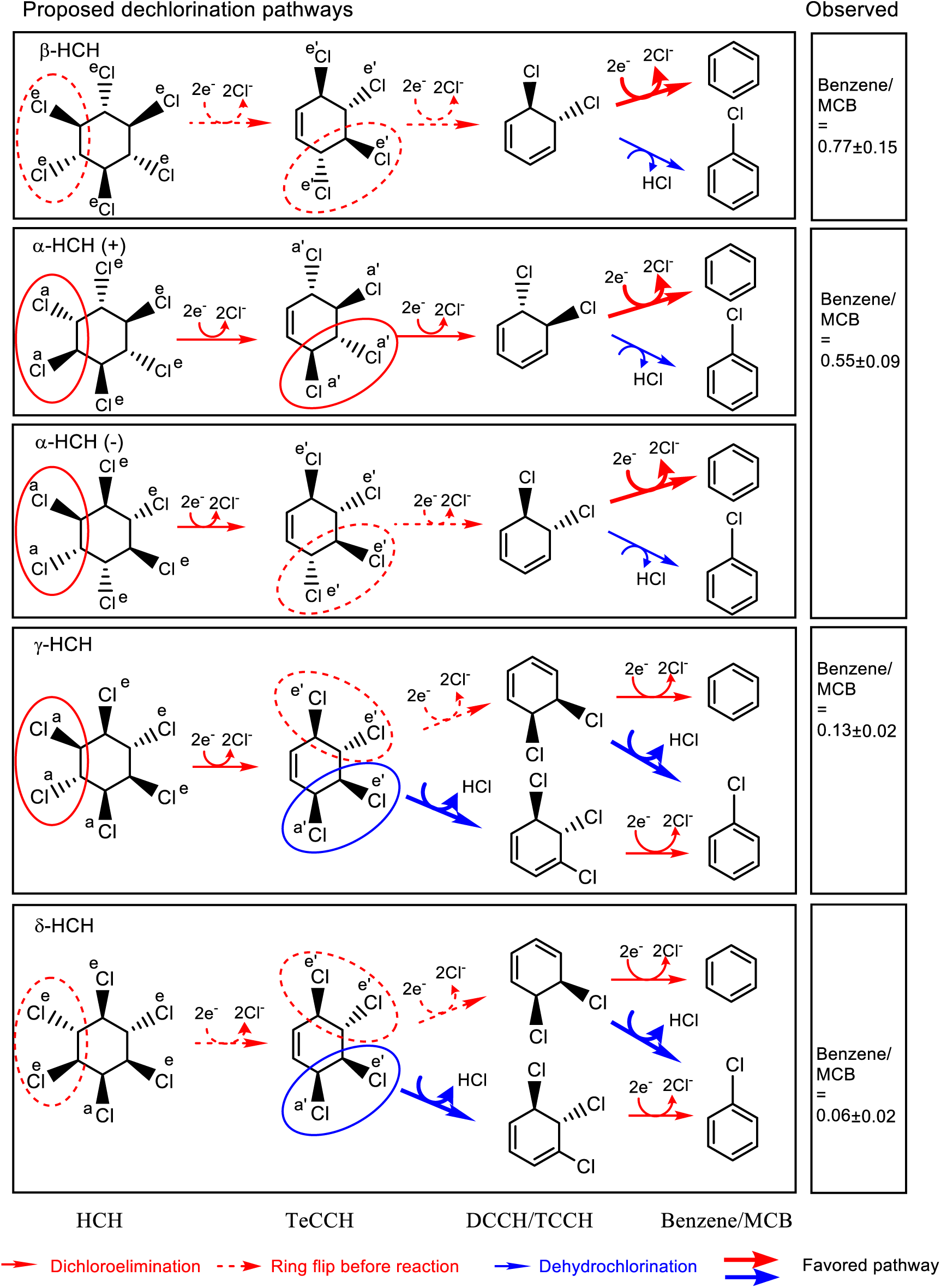
Proposed dechlorination pathways for HCH isomers and observed benzene to MCB molar ratios. Solid wedges indicate bonds that project outward towards the viewer (or “up” on the ring), and dashed wedges are bonds that recede away from the viewer (or “down” on the ring). The superscripts “e” and “a” on the chlorine atoms indicate equatorial or axial positions. Circled substituents highlight leaving groups where the solid lines represent direct elimination while dashed lines mean a ring flip must occur before elimination. Thicker arrows are more favored pathways. Isomers are listed in order of decreasing observed and predicted benzene formation. Figures were generated with ChemDraw (v16).

The theoretical pathway predictions shown in Figure 4 are in accordance with laboratory observations: the dechlorination end-products are benzene and MCB only, and more benzene is produced from the dechlorination of α- and β-HCH than from γ- and δ-HCH. MCB abundance is always greater than benzene in general, however, indicating that the spontaneous dehydrochlorination of DCCH is favored over dichloroelimination. Since we did not observe any dechlorination of MCB to benzene throughout the long incubation period, MCB was not a precursor of benzene in these experiments, and both MCB and benzene were biotransformation end-products of HCH. Please refer to “Dechlorination of HCH Isomers” in the supplementary experimental details for additional discussion on this topic.

### 3.3 Ordination Analysis

Microbial community diversity was surveyed through 16S rRNA gene amplicon sequencing, and the absolute cell numbers of archaea and bacteria were quantified using qPCR. The numbers of quality amplicon sequences obtained per sample are provided in Table S5. The qPCR data and phylogenetic assignments of sequences clustered into OTUs as well as representative sequences for each OTU are provided in Table S6 and S7. An NMDS analysis was performed to visualize the microbial communities and to explore the environmental variables which may relate to the observed differences. Absolute abundances of the most abundant bacteria and archaea (>2%, Table S7F and S7G) were analyzed separately, and the results are shown in Figures 5 and S3. The corresponding stress with 2 dimensions was 0.12 (Figure S4). In Figure 5, the 95% confidence ellipses, representing samples from different isomers, were distinguished from each other, confirming that the type of isomer led to distinct bacterial communities. Within each isomer set, the samples taken on Day 240 clustered separately from those collected on Day 595, indicating the development of microbial communities over time, particularly as donor addition was reduced. Figure 5 also plots the location of bacteria that drive the calculated distances separating the samples. There are some bacteria which are poorly characterized and belong to candidate phyla with unknown cultivated members (unclassified *W22*, WCHB1-15, TA06, OD1, GIF10 and Cloacamonaceae). For example, GIF10, OD1 and Cloacamonaceae are close to samples collected from β-HCH enrichment cultures on Day 595. *Dehalobacter* is located in the middle of the plot closest to the δ-HCH enrichment culture samples. In contrast, an NMDS analysis using the absolute abundances of the most abundant archaea (>2%) revealed only two clusters: one cluster with the parent culture and another cluster comprised of all other culture bottles, regardless of time sampled (Figure S3), indicating that the HCH isomer did not influence archaeal community.

**Figure 5.**
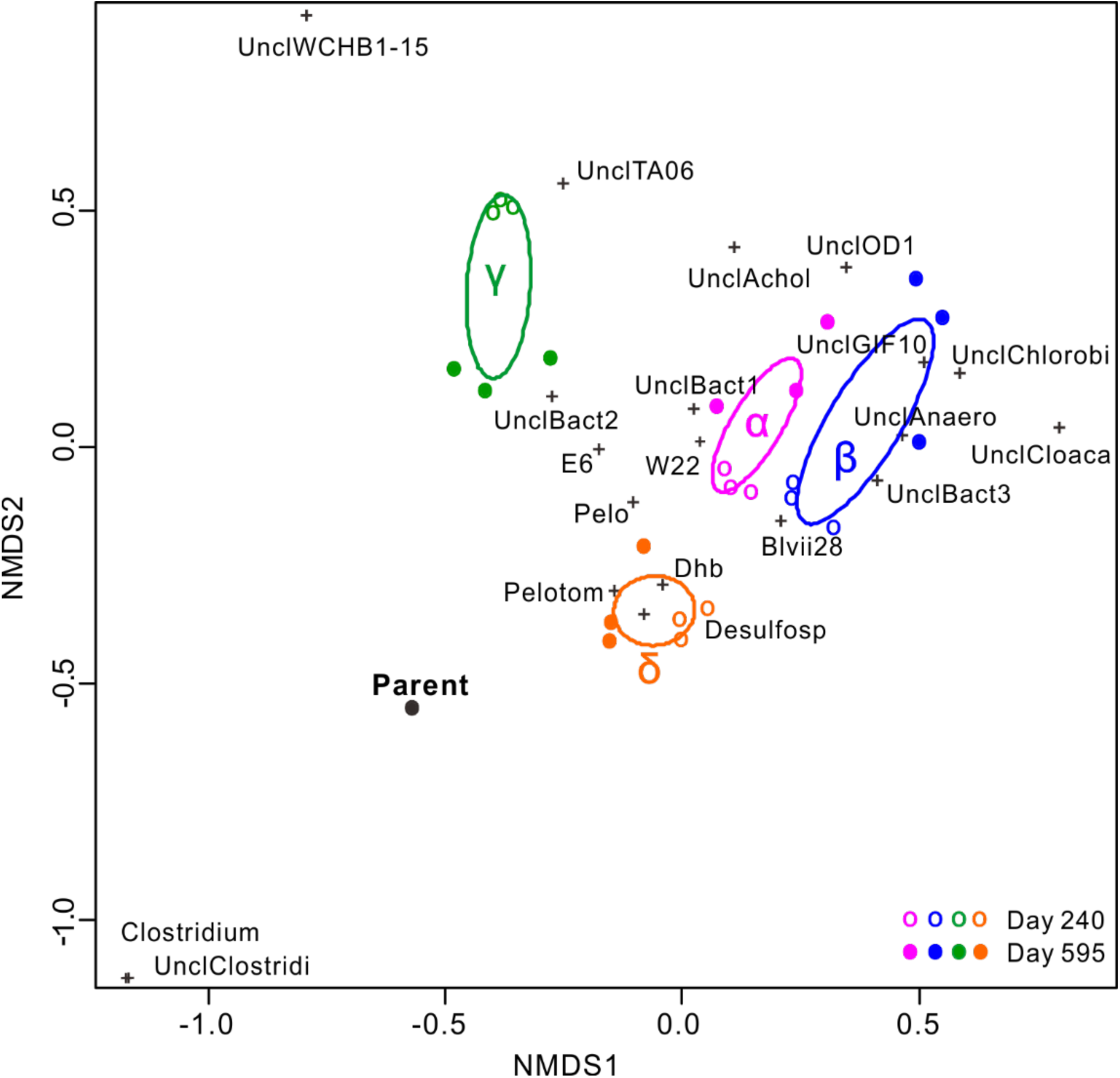
Phylogenetic distances between samples using absolute abundances of top 20 bacteria (>2%, Table S7F), expressed using an NMDS plot (projected from 2 dimensions, stress = 0.12). Distances between points in this figure reflected the rank differences between samples (i.e., dissimilarities). The black, magenta, blue, green and orange points represent the parent GT1, α-, β-, γ- and δ-HCH enrichment cultures, respectively. The open circles are samples collected on Day 240 and closed circles are samples from Day 595. The 95% confidence ellipses for each isomer are also presented. Microbial taxa shown with abbreviated names (Table S7F) were overlaid on the NMDS plot. The archaeal community was not significantly affected by the different HCH isomers (see Figure S3).

### 3.4 Microbial Community Composition

Based on qPCR, a higher fraction of archaea relative to bacteria was observed in enriched samples (Figure S5, Table S6). Given the frequent feedings with excess electron donors, this is not surprising. However, the relative abundance of archaeal groups on Day 595 was not noticeably lower than that of Day 240 even though electron donor addition was reduced and harmonized from Day 499 onwards. The total number of bacteria as inferred from qPCR (Table S6) was on the order of 4×10^7^ 16S rRNA gene copies per mL culture and did not differ much between cultures.

The most abundant (>2%) bacteria and archaea are visualized in a heat map (Figure 6) to facilitate comparisons and to try to identify genera potentially involved in reductive dechlorination of HCH isomers. The heat map shows that triplicate experimental culture bottles yielded similar community profiles, giving assurance that differences between treatments reflect specific amendments. The most abundant taxon in all bottles was affiliated with *Pelobacter* and comprised up to 70% of bacteria in the α- and γ-HCH enrichments towards the end of the study on Day 595. The corresponding representative sequence (501 bp long; Table S7) shares 98.4% sequence identity with *Pelobacter propionicus* DSM 2379. *P. propionicus* was reported to ferment ethanol to acetate and propionate.^62^ The *Pelobacter* enriched in these cultures likely fermented electron donor to provide hydrogen (or acetate) for dechlorination, but is unlikely directly implicated in the dechlorination of the HCH isomers because its abundance decreased when the amount of electron donors added was decreased, specifically in the β-HCH and α-HCH enrichment cultures (Figure 6).

**Figure 6.**
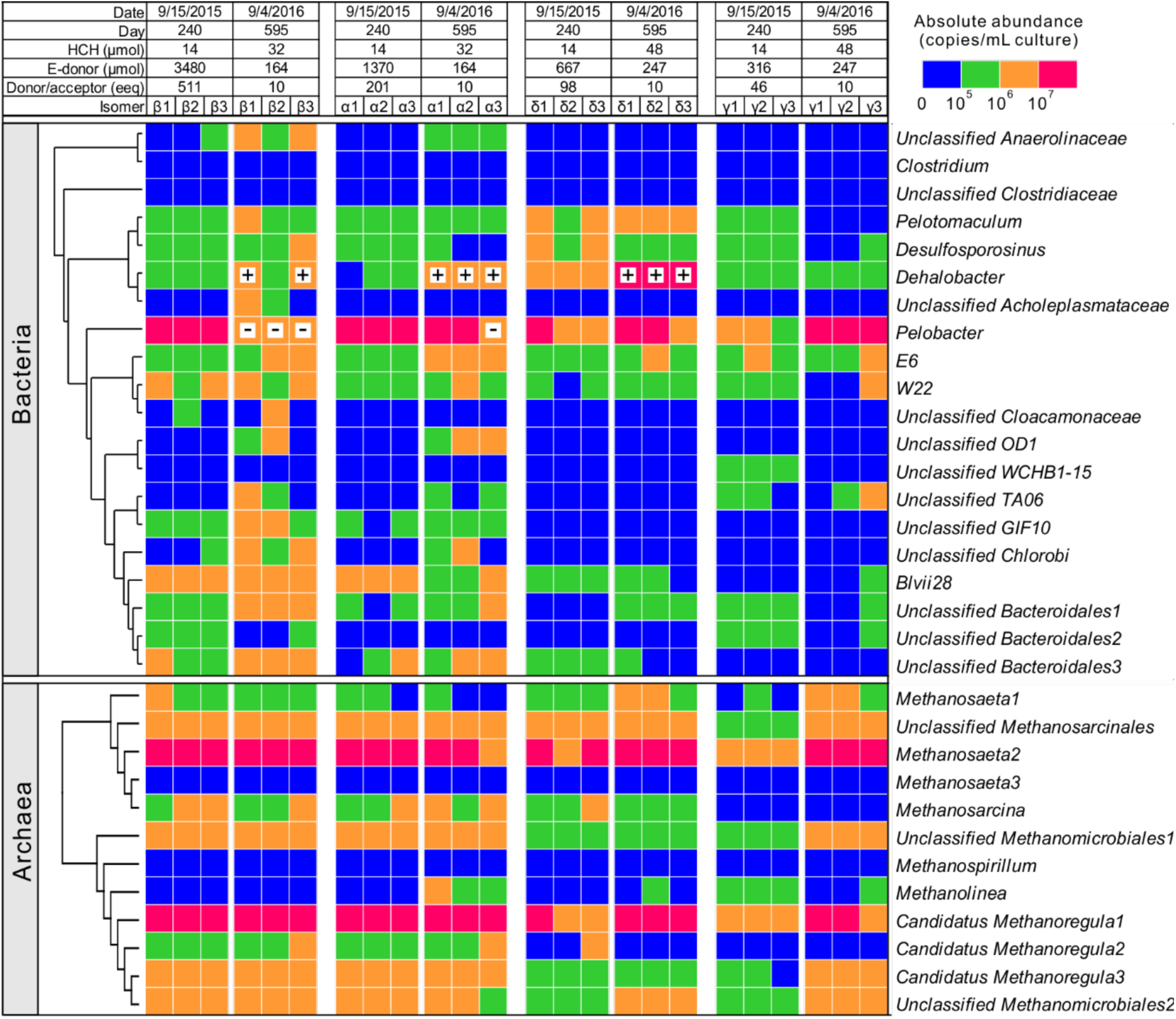
Absolute abundance of major (>2%) bacteria and archaea in enrichment cultures dechlorinating β-, α-, δ- and γ-HCH, listed in order of decreasing amount of cumulative electron donor added. The data above the heat map show DNA sampling date and day, and the total electron donor and HCH amended to each bottle over the 90-day period prior to DNA sampling. In all cases, the amount of donor decreased while the amount of HCH increased between the two time points. At the first sampling time (Day 240), a large excess of donor was provided, particularly for β- and α-isomers. By the second sampling point (Day 595), the donor addition had been harmonized and reduced to10 electron equivalents in excess of that required to dechlorinate added HCH (assuming 6 eeq/mole). The plus signs on the heatmap highlight significant increases of *Dehalobacter* between the two time points, while the minus signs show the decrease of *Pelobacter* in the two culture sets that originally had greatest excess of donor. The phylogenetic tree on the left was created using the PHYML plugin in Geneious (V8.1.9) under the JC69 mode of evolution based on the most abundant bacterial and archaeal OTUs obtained from amplicon sequencing. Taxonomical assignments are on the right. The absolute abundances of each OTU were calculated by multiplying the normalized relative abundances of bacteria or archaea by their total abundance determined by qPCR. The details of the feeding information, qPCR data, absolute/relative abundances of microbes and representative sequences are included in Table S3, S6, S7 and Figure S5.

In contrast, the absolute abundance of *Dehalobacter* increased from Day 240 to Day 595 in all enrichment cultures, making *Dehalobacter* a likely candidate for dechlorinating HCH isomers. *Dehalobacter sp.* E1 was reported to metabolize β-HCH.^19, 44^ To further explore the role of each OTU in these cultures, we calculated the changes in 16S rRNA gene copies per *µ*mol chloride released over the intervals from which we had data and in particular, the last interval from Day 240 to Day 595 (Table S8). Although the time interval was very long, the calculated growth yields of *Dehalobacter* for α-, β- and δ-HCH enrichment cultures ranged from 2×10^5^ to ∼8×10^6^ copies/μmol of chloride released, which is similar to a published growth yield of 1.62×10^6^ copies/μmol for *Dehalobacter* growing on chlorobenzenes.^63, 64^ As *Dehalobacter* is an obligate organohalide-respiring bacterium, these data strongly support that dechlorination of these HCH isomers is metabolic and growth-linked. Only in the γ-HCH enrichment cultures did *Dehalobacter* copies not increase as much from Day 240 to Day 595. Perhaps the fact that these cultures were set up 130 days later, and were provided with less electron donors overall affected these results. The only other OTU in the γ-HCH enrichments with a significant increase between days 240 and 595 is *Pelobacter* (Table S8D), yet this OTU is unlikely an organohalide-respiring bacterium, as reasoned above. Other OTUs that showed a consistent pattern of increasing abundance and yield for a specific isomer between Day 240 and Day 595 (Figure 6 and Table S8) include unclassified *Synergistales*_E6 (α-, β- and δ-HCH), unclassified OD1 (α-HCH), unclassified *Chlorobi* and unclassified GIF10 (β-HCH), and unclassified *Bacteroidales* (α- and β-HCH). Further work is clearly needed to determine if any of these poorly characterized organisms are responsible for dechlorination, although most seem likely to be fermenting organisms or biomass recyclers. The dominant archaea, *Methanosaeta* and *Candidatus Methanoregula* (Figure 6) are known to use fermentation products such as acetate and hydrogen to produce methane (shown in Figure 1).

### 3.5 Implications for HCH contaminated sites

HCH contamination is extensive on former agricultural lands as well as in legacy production and waste storage or containment facilities around the world. The stable HCH-dechlorinating microbial enrichments cultures obtained in this study are important building blocks in the development of bioremediation technologies for HCH-contaminated sites. In fact, a successful demonstration of HCH biodegradation (via biostimulation) in a full scale *in situ* bioscreen was reported in the Netherlands.^65^ Such approaches would benefit from the added value of *in situ* bioaugmentation. Though partial dechlorination to MCB and benzene is not in itself a full remediation, a combination of technologies could be used to achieve complete bioconversion of HCH to non-toxic end products, depending on site conditions. Moreover, benzene and MCB sorb less than the parent compounds and thus can be ultimately removed from soil more easily. Bioaugmentation with different cultures capable of biotransformation of MCB and benzene may be feasible. In fact, the anaerobic conversion of benzene and MCB in bioaugmented microcosms has been reported.^66^ A combination of anaerobic and aerobic treatments may also be feasible to take advantage of the fast aerobic biodegradation of MCB and benzene. Further investigations into the mechanism of dechlorination and the feasibility of complete detoxification of HCH isomers using multiple enrichment cultures are underway.

## AUTHOR CONTRIBUTIONS

E.A.E., S.G., X.T., and L.A.P.J. conceived the work. W.Q., L.A.P.J., I.R.A., and X.T. performed the experimental work at BioZone, University of Toronto. W.Q., L.A.P.J, and E.A.E wrote the manuscript. W.Q. and I.Y. performed the bioinformatics analyses. All authors analyzed the data, reviewed and provided comments on the manuscript.

## ASSOCIATED CONTENT

### Supporting Information

The Supplemental files contain Figures S1-S5, Table S1-S8, and method details. Included is information on the stereochemistry, basic physical-chemical and thermodynamic properties of HCH isomers, feeding information, 16S amplicon sequencing data, NMDS analysis with archaeal sequencing data, qPCR raw data, and yield calculations. Additional information for method details includes substrate amendment methods, qPCR, amplicon sequencing and dechlorination mechanisms.

### Notes

The authors declare no competing financial interest.

## Acknowledgement

We gratefully acknowledge funding from the Key Program for International S&T Cooperation Projects of China (Ontario-China Research and Innovation Fund, 2016YFE0101900), the Government of Canada through Genome Canada and the Ontario Genomics Institute (OGI-102). W.Q. and X. T. were supported by the China Scholarship Council. L. P. J. was supported by the NSERC CREATE RENEW program [180804567], the Government of Ontario through the Ontario Graduate Scholarship program and the Ontario Research Fund INTEGRATE project [ORF-RE05-WR-01].

## REFERENCES

1. Kutz, F. W.; Wood, P. H.; Bottimore, D. P., Organochlorine pesticides and polychlorinated biphenyls in human adipose tissue. Rev. Environ. Contam. Toxicol. 1991, 120, (6), 1–82.

2. Dominguez, C. M.; Rodriguez, S.; Lorenzo, D.; Romero, A.; Santos, A., Degradation of hexachlorocyclohexanes (HCHs) by stable zero valent iron (ZVI) microparticles. Water Air Soil Pollut. 2016, 227, (12), 446.

3. Willett, K. L.; Ulrich, E. M.; Hites, R. A., Differential toxicity and environmental fates of hexachlorocyclohexane isomers. Environ. Sci. Technol. 1998, 32, (15), 2197–2207.

4. Liu, Y.; Bashir, S.; Stollberg, R.; Trabitzsch, R.; Weiß, H.; Paschke, H.; Nijenhuis, I.; Richnow, H.-H., Compound specific and enantioselective stable Isotope analysis as tools to monitor transformation of hexachlorocyclohexane (HCH) in a complex aquifer system. Environ. Sci. Technol. 2017, 51, (16), 8909–8916.

5. Middeldorp, P. J.; Jaspers, M.; Zehnder, A. J.; Schraa, G., Biotransformation of α-, ß-, γ-, and δ-hexachlorocyclohexane under methanogenic conditions. Environ. Sci. Technol. 1996, 30, (7), 2345–2349.

6. Lal, R.; Pandey, G.; Sharma, P.; Kumari, K.; Malhotra, S.; Pandey, R.; Raina, V.; Kohler, H.-P. E.; Holliger, C.; Jackson, C., Biochemistry of microbial degradation of hexachlorocyclohexane and prospects for bioremediation. Microbiol. Mol. Biol. Rev. 2010, 74, (1), 58–80.

7. Alvarez, A.; Benimeli, C. S.; Saez, J. M.; Fuentes, M. S.; Cuozzo, S. A.; Polti, M. A.; Amoroso, M. J., Bacterial bio-resources for remediation of hexachlorocyclohexane. Int. J. Mol. Sci. 2012, 13, (11), 15086–106.

8. Saez, J.; Alvarez, A.; Fuentes, M.; Amoroso, M.; Benimeli, C., An Overview on Microbial Degradation of Lindane. In Microbe-Induced Degradation of Pesticides, Springer: 2017; pp 191–212.

9. Phillips, T. M.; Seech, A. G.; Lee, H.; Trevors, J. T., Biodegradation of hexachlorocyclohexane (HCH) by microorganisms. Biodegradation 2005, 16, (4), 363–392.

10. Mehboob, F.; Langenhoff, A. A.; Schraa, G.; Stams, A. J., Anaerobic degradation of lindane and other HCH Isomers. In Management of Microbial Resources in the Environment, Springer: 2013; pp 495–521.

11. Bashir, S.; Hitzfeld, K. L.; Gehre, M.; Richnow, H. H.; Fischer, A., Evaluating degradation of hexachlorcyclohexane (HCH) isomers within a contaminated aquifer using compound-specific stable carbon isotope analysis (CSIA). Water Res. 2015, 71, 187–196.

12. Lian, S.; Nikolausz, M.; Nijenhuis, I.; Francisco Leite, A.; Richnow, H. H., Biotransformation and inhibition effects of hexachlorocyclohexanes during biogas production from contaminated biomass characterized by isotope fractionation concepts. Bioresource Technol. 2018, 250, 683–690.

13. Mackay, D.; Shiu, W. Y.; Ma, K.-C.; Lee, S. C., Physical-chemical properties and environmental fate for organic chemicals. 2nd ed.; CRC Press: 2006; Vol. 5.

14. Clayton, G. D.; Clayton, F. E., Patty’s industrial hygiene and toxicology. 3rd ed.; John Wiley & Sons: 1981.

15. Bruice, P. Y., Organic Chemistry. Pearson Education, Inc.: United States of America, 2017.

16. Badea, S.-L.; Vogt, C.; Weber, S.; Danet, A.-F.; Richnow, H.-H., Stable isotope fractionation of gamma-hexachlorocyclohexane (lindane) during reductive dechlorination by two strains of sulfate-reducing bacteria. Environ. Sci. Technol. 2009, 43, (9), 3155.

17. Li, S.; Elliott, D. W.; Spear, S. T.; Ma, L.; Zhang, W.-X., Hexachlorocyclohexanes in the Environment: Mechanisms of Dechlorination. Crit. Rev. Env. Sci. Tec. 2011, 41, (19), 1747–1792.

18. Buser, H.-R.; Mueller, M. D., Isomer and enantioselective degradation of hexachlorocyclohexane isomers in sewage sludge under anaerobic conditions. Environ. Sci. Technol. 1995, 29, (3), 664–672.

19. Doesburg, W.; Eekert, M. H.; Middeldorp, P. J.; Balk, M.; Schraa, G.; Stams, A. J., Reductive dechlorination of ß-hexachlorocyclohexane (ß-HCH) by a Dehalobacter species in coculture with a Sedimentibacter sp. FEMS Microbiol. Ecol. 2005, 54, (1), 87–95.

20. Cristol, S. J., The kinetics of the alkaline dehydrochlorination of the benzene hexachloride Isomers. The mechanism of second-order elimination reactions. J. Am. Chem. Soc. 1947, 69, (2), 338–342.

21. Liu, X.; Peng, P. a.; Fu, J.; Huang, W., Effects of FeS on the transformation kinetics of γ-hexachlorocyclohexane. Environ. Sci. Technol. 2003, 37, (9), 1822–1828.

22. Mackenzie, K.; Battke, J.; Koehler, R.; Kopinke, F.-D., Catalytic effects of activated carbon on hydrolysis reactions of chlorinated organic compounds: Part 2. 1,1,2,2- Tetrachloroethane. Appl. Catal. B-Environ. 2005, 59, (3), 171–179.

23. Beland, F. A.; Farwell, S. O.; Robocker, A. E.; Geer, R. D., Electrochemical reduction and anaerobic degradation of lindane. J. Agr. Food Chem. 1976, 24, (4), 753.

24. Birkin, P. R.; Evans, A.; Milhano, C.; Montenegro, M. I.; Pletcher, D., The Mediated Reduction of Lindane in DMF. Electroanalysis 2004, 16, (7), 583–587.

25. Hughes, E. D.; Ingold, C. K.; Pasternak, R., Mechanism of elimination reactions. Part XVIII. Kinetics and steric course of elimination from isomeric benzene hexachlorides. J. Chem. Soc. 1953, 3832–3839.

26. Bhatt, P.; Kumar, M. S.; Chakrabarti, T., Fate and degradation of POP- hexachlorocyclohexane. Crit. Rev. Env. Sci. Tec. 2009, 39, (8), 655–695.

27. Lal, R.; Dogra, C.; Malhotra, S.; Sharma, P.; Pal, R., Diversity, distribution and divergence of lin genes in hexachlorocyclohexane-degrading sphingomonads. Trends Biotechnol. 2006, 24, (3), 121–130.

28. Bashir, S.; Fischer, A.; Nijenhuis, I.; Richnow, H.-H., Enantioselective Carbon Stable Isotope Fractionation of Hexachlorocyclohexane during Aerobic Biodegradation by Sphingobium spp. Environ. Sci. Technol. 2013, 47, (20), 11432–11439.

29. Tu, C. M., Utilization and degradation of lindane by soil microorganisms. Arch. Microbiol. 1976, 108, (3), 259–263.

30. Boltner, D.; Moreno-Morillas, S.; Ramos, J. L., 16S rDNA phylogeny and distribution of lin genes in novel hexachlorocyclohexane-degrading Sphingomonas strains. Environ. Microbiol. 2005, 7, (9), 1329–1338.

31. Senoo, K.; Wada, H., Isolation and identification of an aerobic gamma-HCH decomposing bacterium from soil. Soil Sci. Plant Nutr. 1989, 35, (1), 79–87.

32. Dogra, C.; Raina, V.; Pal, R.; Suar, M.; Lal, S.; Gartemann, K. H.; Holliger, C.; van der Meer, J. R.; Lal, R., Organization of lin genes and IS6100 among different strains of hexachlorocyclohexane-degrading Sphingomonas paucimobilis: Evidence for horizontal gene transfer. J. Bacteriol. 2004, 186, (8), 2225–2235.

33. Kumari, R.; Subudhi, S.; Suar, M.; Dhingra, G.; Raina, V.; Dogra, C.; Lal, S.; van der Meer, J. R.; Holliger, C.; Lal, R., Cloning and characterization of lin genes responsible for the degradation of hexachlorocyclohexane isomers by Sphingomonas paucimobilis strain B90. Appl. Environ. Microb. 2002, 68, (12), 6021–6028.

34. Vijgen, J.; Weber, R.; Lichtensteiger, W.; Schlumpf, M., The legacy of pesticides and POPs stockpiles-a threat to health and the environment. Environ. Sci. Pollut. Res. Int. 2018, 25, (32), 31793–31798.

35. Laquitaine, L.; Durimel, A.; de Alencastro, L. F.; Jean-Marius, C.; Gros, O.; Gaspard, S., Biodegradability of HCH in agricultural soils from Guadeloupe (French West Indies): identification of the lin genes involved in the HCH degradation pathway. Environ. Sci. Pollut. R. 2016, 23, (1), 120–127.

36. Middeldorp, P. J.; van Doesburg, W.; Schraa, G.; Stams, A. J., Reductive dechlorination of hexachlorocyclohexane (HCH) isomers in soil under anaerobic conditions. Biodegradation 2005, 16, (3), 283–290.

37. Tsukano, Y.; Kobayashi, A., Formation of γ-BTC in flooded rice field soils treated with γ-BHC. Agr. Biol. Chem. 1972, 36, 166–167.

38. Rodríguez-Garrido, B.; Lú-Chau, T. A.; Feijoo, G.; Macías, F.; Monterrroso, M. C., Reductive Dechlorination of α-, ß-, γ-, and δ-Hexachlorocyclohexane Isomers with Hydroxocobalamin, in Soil Slurry Systems. Environ. Sci. Technol. 2010, 44, (18), 7063.

39. MacRae, I.; Raghu, K.; Bautista, E., Anaerobic degradation of the insecticide lindane by Clostridium sp. Nature 1969, 221, (5183), 859–860.

40. Jagnow, G.; Haider, K.; Ellwardt, P.-C., Anaerobic dechlorination and degradation of hexachlorocyclohexane isomers by anaerobic and facultative anaerobic bacteria. Arch. Microbiol. 1977, 115, (3), 285–292.

41. Ohisa, N.; Yamaguchi, M., Gamma BHC degradation accompanied by the growth of Clostridium rectum isolated from paddy field soil. Agric. Biol. Chem. 1978, 42, (10), 1819–1823.

42. Boyle, A. W.; Häggblom, M. M.; Young, L. Y., Dehalogenation of lindane (γ- hexachlorocyclohexane) by anaerobic bacteria from marine sediments and by sulfate-reducing bacteria. FEMS Microbiol. Ecol. 1999, 29, (4), 379–387.

43. Elango, V.; Kurtz, H. D.; Anderson, C.; Freedman, D. L., Use of δ- hexachlorocyclohexane as a terminal electron acceptor by an anaerobic enrichment culture. J. Hazard. Mater. 2011, 197, 204–210.

44. Maphosa, F.; van Passel, M. W. J.; de Vos, W. M.; Smidt, H., Metagenome analysis reveals yet unexplored reductive dechlorinating potential of Dehalobacter sp. E1 growing in co-culture with Sedimentibacter sp. Env. Microbiol. Rep. 2012, 4, (6), 604–616.

45. Bashir, S.; Kuntze, K.; Vogt, C.; Nijenhuis, I., Anaerobic biotransformation of hexachlorocyclohexane isomers by Dehalococcoides species and an enrichment culture. Biodegradation 2018, 29, (4), 409–418.

46. Baker, M. T.; Nelson, R. M.; Van Dyke, R. A., The formation of chlorobenzene and benzene by the reductive metabolism of lindane in rat liver microsomes. Arch. Biochem. Biophys. 1985, 236, (2), 506–514.

47. Heritage, A.; Rae, I., Identification of intermediates formed during the degradation of hexachlorocyclohexanes by Clostridium sphenoides. Appl. Environ. Microbiol. 1977, 33, (6), 1295–1297.

48. Edwards, E. A.; Grbic-Galic, D., Anaerobic degradation of toluene and o-xylene by a methanogenic consortium. Appl. Environ. Microbiol. 1994, 60, (1), 313–322.

49. Amann, R. I.; Ludwig, W.; Schleifer, K.-H., Phylogenetic identification and in situ detection of individual microbial cells without cultivation. Microbiol. Rev. 1995, 59, (1), 143–169.

50. Ferris, M. J.; Muyzer, G.; Ward, D. M., Denaturing gradient gel electrophoresis profiles of 16S rRNA-defined populations inhabiting a hot spring microbial mat community. Appl. Environ. Microbiol. 1996, 62, (2), 340.

51. Engelbrektson, A.; Kunin, V.; Wrighton, K. C.; Zvenigorodsky, N.; Chen, F.; Ochman, H.; Hugenholtz, P., Experimental factors affecting PCR-based estimates of microbial species richness and evenness. ISME J. 2010, 4, (5), 642–647.

52. Hanshew, A. S.; Mason, C. J.; Raffa, K. F.; Currie, C. R., Minimization of chloroplast contamination in 16S rRNA gene pyrosequencing of insect herbivore bacterial communities. J. Microbiol. Methods 2013, 95, (2), 149–155.

53. Ferris, M.; Muyzer, G.; Ward, D., Denaturing gradient gel electrophoresis profiles of 16S rRNA-defined populations inhabiting a hot spring microbial mat community. Appl. Environ. Microbiol. 1996, 62, (2), 340–346.

54. Caporaso, J. G.; Kuczynski, J.; Stombaugh, J.; Bittinger, K.; Bushman, F. D.; Costello, E. K.; Fierer, N.; Pena, A. G.; Goodrich, J. K.; Gordon, J. I.; Huttley, G. A.; Kelley, S. T.; Knights, D.; Koenig, J. E.; Ley, R. E.; Lozupone, C. A.; McDonald, D.; Muegge, B. D.; Pirrung, M.; Reeder, J.; Sevinsky, J. R.; Turnbaugh, P. J.; Walters, W. A.; Widmann, J.; Yatsunenko, T.; Zaneveld, J.; Knight, R., QIIME allows analysis of high-throughput community sequencing data. Nat. Methods 2010, 7, (5), 335–336.

55. Edgar, R. C., Search and clustering orders of magnitude faster than BLAST. Bioinformatics 2010, 26, (19), 2460–2461.

56. Haas, B. J.; Gevers, D.; Earl, A. M.; Feldgarden, M.; Ward, D. V.; Giannoukos, G.; Ciulla, D.; Tabbaa, D.; Highlander, S. K.; Sodergren, E., Chimeric 16S rRNA sequence formation and detection in Sanger and 454-pyrosequenced PCR amplicons. Genome Res. 2011, 21, (3), 494.

57. Cole, J. R.; Wang, Q.; Fish, J. A.; Chai, B.; McGarrell, D. M.; Sun, Y.; Brown, C. T.; Porras-Alfaro, A.; Kuske, C. R.; Tiedje, J. M., Ribosomal Database Project: data and tools for high throughput rRNA analysis. Nucleic Acids Res. 2014, 42, (D1), D633–D642.

58. McDonald, D.; Price, M. N.; Goodrich, J.; Nawrocki, E. P.; DeSantis, T. Z.; Probst, A.; Andersen, G. L.; Knight, R.; Hugenholtz, P., An improved Greengenes taxonomy with explicit ranks for ecological and evolutionary analyses of bacteria and archaea. ISME J. 2012, 6, (3), 610–618.

59. Clarke, K. R., Non-parametric multivariate analyses of changes in community structure. Aust. J. Ecol. 1993, 18, (1), 117–143.

60. Kocur, C. M.; Lomheim, L.; Molenda, O.; Weber, K. P.; Austrins, L. M.; Sleep, B. E.; Boparai, H. K.; Edwards, E. A.; O’Carroll, D. M., Long-term field study of microbial community and dechlorinating activity following carboxymethyl cellulose-stabilized nanoscale zero-valent iron injection. Environ. Sci. Technol. 2016, 50, (14), 7658–7670.

61. Ohisa, N.; Yamaguchi, M.; Kurihara, N., Lindane degradation by cell-free extracts of Clostridium rectum. Arch. Microbiol. 1980, 125, (3), 221–225.

62. Schink, B., Fermentation of 2,3-butanediol by Pelobacter carbinolicus sp. nov. and Pelobacter propionicus sp. nov., and evidence for propionate formation from compounds. Arch. Microbiol. 1984, 137, (1), 33–41.

63. Nelson, J. L.; Jiang, J.; Zinder, S. H., Dehalogenation of chlorobenzenes, dichlorotoluenes, and tetrachloroethene by three Dehalobacter spp. Environ. Sci. Technol. 2014, 48, (7), 3776–3782.

64. Puentes Jácome, L. A.; Edwards, E. A., A switch of chlorinated substrate causes emergence of a previously undetected native Dehalobacter population in an established Dehalococcoides-dominated chloroethene-dechlorinating enrichment culture. FEMS Microbiol. Ecol. 2017, 93, (12).

65. Langenhoff, A. A.; Staps, S. J.; Pijls, C.; Rijnaarts, H. H., Stimulation of hexachlorocyclohexane (HCH) biodegradation in a full scale in situ bioscreen. Environ. Sci. Technol. 2013, 47, (19), 11182–8.

66. Liang, X.; Devine, C. E.; Nelson, J.; Lollar, B. S.; Zinder, S.; Edwards, E. A., Anaerobic conversion of chlorobenzene and benzene to CH4 and CO2 in bioaugmented Microcosms. Environ. Sci. Technol. 2013, 47, (5), 2378–2385.

