## Supplemental information text file for "Microbial Communities Associated with Sustained Anaerobic Reductive Dechlorination of α-, β-, γ-, and δ-Hexachlorocyclohexane Isomers to Monochlorobenzene and Benzene"

Number of Pages: 15
Number of Figures: 5
Number of Tables: 8
*Supplementary Tables S3, S6, S7 and S8 are included in an Excel file separately.

### TABLE OF CONTENTS


**List of Supporting Figures**

**Figure S1.** Stereochemistry of α-, β-, γ-, and δ-HCH investigated in this study.

**Figure S2**. Concentration profiles of benzene and monochlorobenzene (MCB) in the sterile controls.

**Figure S3**. Phylogenetic distances between samples using absolute abundance of archaea (>2%, Table S7G), visualized using an NMDS plot.

**Figure S4.** Shepard stress plot (k=2, stress=0.12).

**Figure S5**. Comparison of relative abundances determined by two methods: amplicon sequencing and qPCR.

**Supplemental Experimental Details**

i. HCH and Electron Donor Amendment Methods
ii. Quantitative PCR
iii. Amplicon Sequencing
iv. Dechlorination of HCH Isomers

**List of Supporting Tables**

**Table S1.** Selected physical and chemical properties of HCH isomers at 25°C.

**Table S2.** Basic thermodynamic information on two conformers of HCH isomers.

**Table S3 (Excel)** Feeding information

**Table S4.** Comparison of the ratios of Benzene to MCB with previous published values for the HCH isomers.

**Table S5.** Number of sequences per sample during data processing

**Table S6 (Excel).** qPCR reactions

**Table S7 (Excel).** Bacteria and Archaea identified in samples (normalized relative and absolute abundance calculations; a series of 7 smaller related tables)

**Table S8 (Excel).** Yield calculations

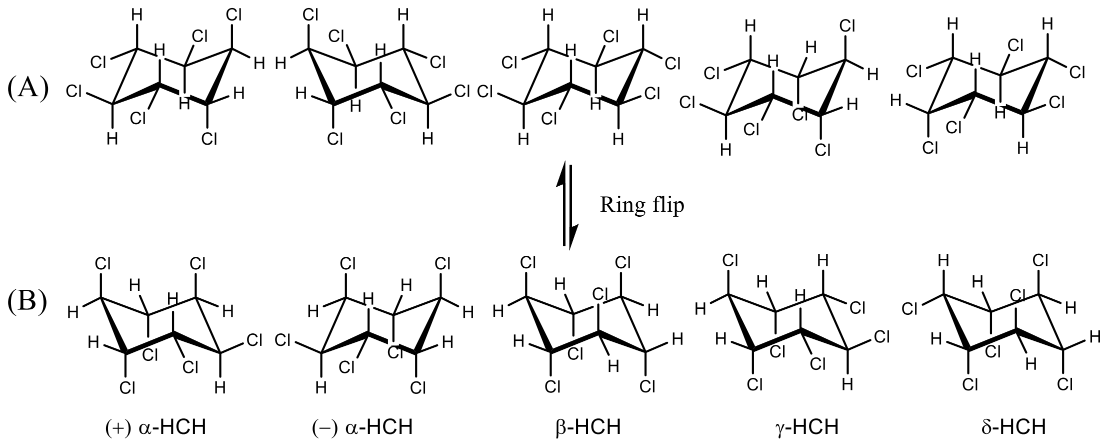


**Figure S1.** Stereochemistry of α-, β-, γ-, and δ-HCH investigated in this study including the two enantiomers of α-HCH. The axial chlorine-carbon bonds are vertical and alternate above and below the ring, while the equatorial chlorine-carbon bonds point outward from the ring. Panel A: thermodynamically stable conformers and the positions of chlorines on the cyclohexane rings are: α: aaeeee, β: eeeeee, γ: aaaeee, δ: aeeeee, (‘a’ represents chlorines on the axial position and ‘e’ represents chlorines on the equatorial position). Panel B: HCH conformer after ring flip in which all axial atoms convert into equatorial and vice versa. Figures were generated in ChemDraw v16.0.1.4 (77).

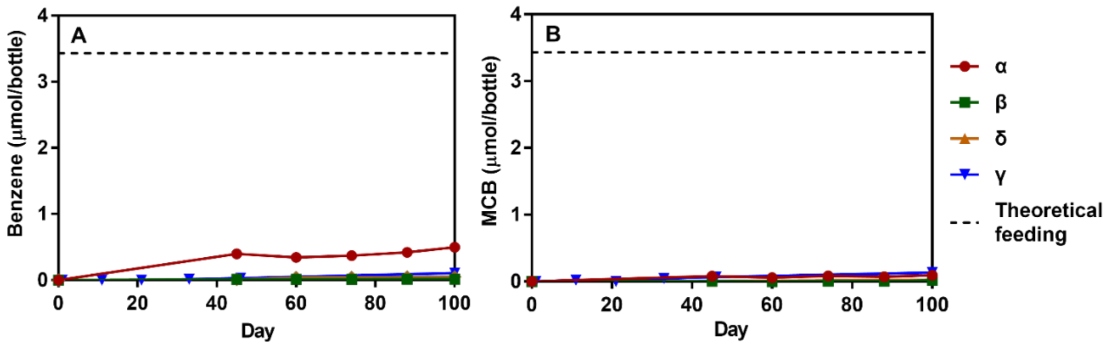


**Figure S2**. Concentration profiles of benzene (panel A) and monochlorobenzene (MCB, panel B) in the sterile controls with autoclaved cultures. The left panel represents the change of benzene, and the right panel shows the change of MCB over 100 days of incubation. Only one initial dose of HCH was provided to these bottles, and of that, 17%, 7%, 2% and 1% of the added α-, γ-, δ- and β-HCH, respectively, was abiotically dechlorinated to benzene and MCB over 100 days of incubation. Only one bottle was prepared for each isomer.

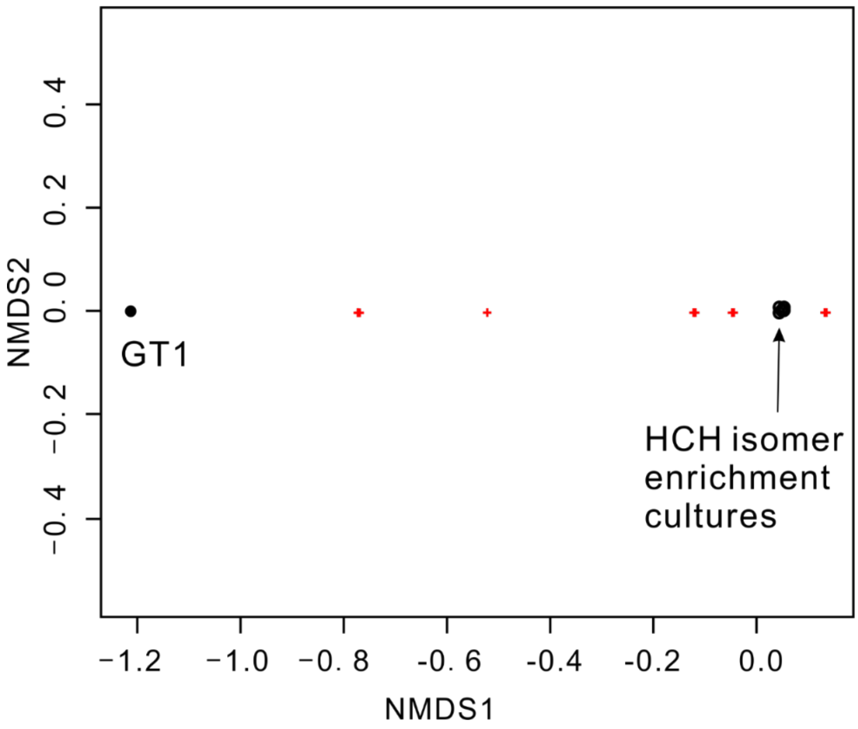


**Figure S3**. Phylogenetic distances between samples (black circles) based on absolute abundance of archaea visualized in an NMDS plot (projected from 2 dimensions, stress of $5\times10^{-5}$). Only two clusters (one cluster with the parent culture (GT1) and another cluster with all other bottles (and all time points) are observed, suggesting that the type of isomer amended, and electron donor changes did not significantly influence the archaeal community. The red "+" signs represent the position of archaea >2% shown in Table S7G.

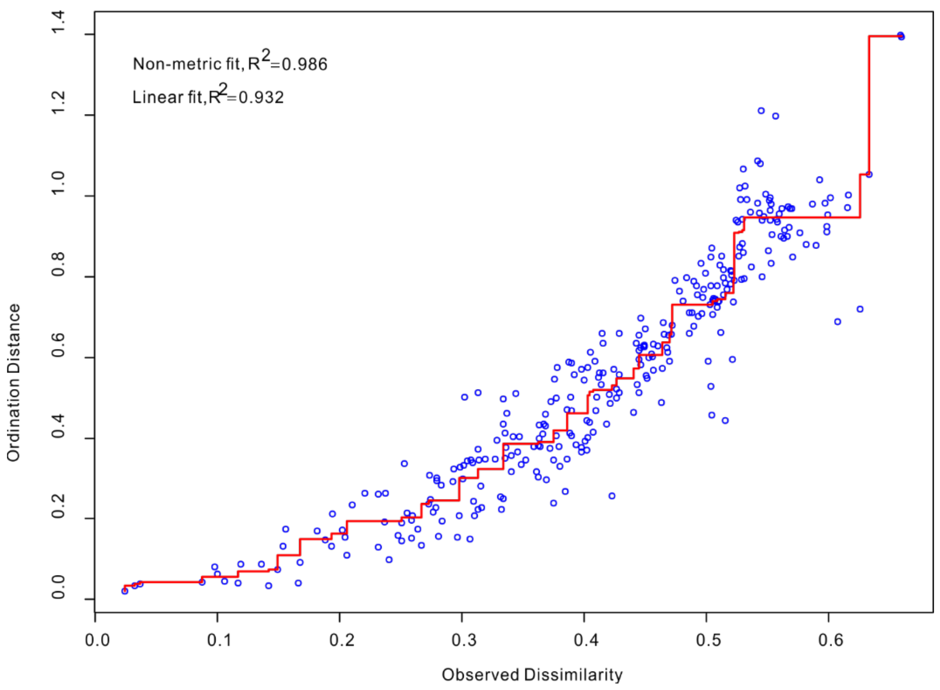


**Figure S4.** Shepard stress plot showing the relationship between the rank order of the actual dissimilarities (from original dissimilarity matrix) and the ordination distances (distances on final plot) at k=2 (the number of dimensions chosen for final model). Stress is a measure of the agreement between ordination distances and calculated dissimilarity scores. If there is a good match and stress is low, all points lie on a steadily increasing line. K=2, stress=0.12

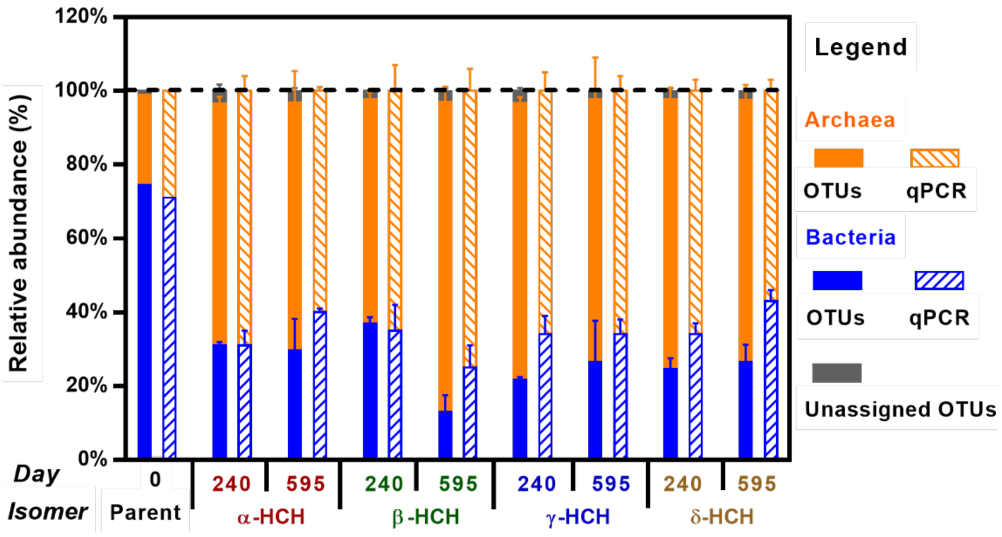


**Figure S5**. Comparison of relative abundances determined by two methods: amplicon sequencing (solid bars) and qPCR (striped bars). Shown results are relative abundances of total bacteria (blue bars), archaea (orange bars) and unassigned OTUs (grey bars, barely visible). Horizontal axis shows sampling time (Day) and isomer. Data on Day 240 and 595 are average of triplicate bottles and error bars represent one standard deviation. qPCR consistently gave higher fraction of bacteria than amplicon sequencing. All primers are subject to some bias as illustrated here.

**Supplemental Experimental Details**

#### i. HCH and Electron Donor Amendment Methods

The solubility of each HCH isomer in water and in ethanol are different (Table S1) because of the different geometries of the cyclohexane ring (isomers shown in Figure S1). As a result, the initial amount of electron donor (ethanol/acetone) amended to each isomer-dechlorinating culture was different. A new feeding method was devised to try to reduce the amount of solvent (donor) added to each microcosm to decrease the proportion of donor-fermenting organisms and to decrease methane production. The new feeding method used HCH isomers dissolved in acetone. The HCH/acetone solution was spiked into clean and autoclaved 250-mL bottles to deliver equal amounts of HCH to each bottle. The bottles were then purged in a fume hood with sterile $N_2$ to evaporate all the acetone. Then the bottles with dry HCH were moved into glovebox and allowed to sit for one week. Finally, 10 mL of fresh medium with redox indicator (resazurin) was added to the new bottles to make sure the bottles were anaerobic (i.e. medium was not pink).

When an enrichment culture was ready to be refed with HCH, the culture bottle was removed from the glove box and purged with a gas mix ($N_2/CO_2$: 80%/20% by volume) to remove the metabolic products (MCB and benzene). The bottles were returned to the glovebox and the entire contents of the cultures (~ 90 mL) was transferred into the new bottles containing solid HCH and 10 mL fresh medium. These new culture bottles with HCH isomers and culture (~100 mL) were purged with $N_2/CO_2$ gas mix to remove hydrogen (present in glove box atmosphere). Ethanol (electron donor; 12 eeq/mol) was amended to each culture at 10 times electron equivalents required for complete dechlorination of HCH (6 eeq/mol). The amounts of HCH isomers and electron donor amended to each culture are provided in Table S3.

#### ii. Quantitative PCR (qPCR)

Each 20 $\mu$L qPCR reaction was prepared in duplicate and dissolved in sterile UltraPure distilled water containing 10 $\mu$L of EvaGreen Supermix (Bio-Rad Laboratories, Herculed, CA), 1 $\mu$L of each 10 $\mu$M forward and reverse primers (described below), 2 $\mu$L of DNA template (or plasmid dilutions). The thermocycling program was as follows: initial denaturation at 98℃ for 2min, followed by 40 cycles of denaturation at 98℃ for 5s, annealing at 55℃ (total bacteria) or 59℃ (total archaea) for 10s and chain extension at 65℃ for 10s.
The primers used for the enumeration of total bacteria (Amann et al. 1995) were: 1055f: 5;-ATGGCTGTCGTCAGCT-3' and 1392r 5'-ACGGGCGGTGTGTAC-3'. The primers used for the amplification of total archaea (Ferris et al. 1996b) were 787f 5'-ATTAGATACCCGBGTAGTCC-3' and 1059r 5'-GCCATGCACCWCCTCT-3'.

#### iii. Amplicon Sequencing

For pyrotag sequencing, the hypervariable V6 to V8 region of the 16S rRNA gene of bacteria, archaea and the 18S rRNA gene in eukaryote was amplified using PCR reactions with the primer set 926f (5'-AAA CTY AAA KGA ATT GAC GG-3') and

1392r (5'-ACG GGC GGT GTG TRC-3') (Engelbrektson et al. 2010; Hanshew et al. 2013) on a MJ Research PTC-200 Peltier Thermal Cycler (Bio-Rad Laboratories, Hercules, CA). The thermocycling program is as follows: 95 ℃, 3min; 25 cycles of 90 ℃ 30s, 54℃ 45s, 72℃ 10min; final hold at 4℃ (Kocur et al. 2016). The forward and reverse primers also included 30 bp adaptors sequences (CCA TCT CAT CCC TGC GTG TCT CCG ACT CAG; CCT ATC CCC TGT GTG CCT TGG CAG TCT CAG, respectively). The reverse primer also contained a 10bp multiplex identifier (MID) bar code to distinguish multiple samples pooled within one sequencing region. The PCR products were verified on a 2% agarose gel and replicates were combined and purified suing GeneJET PCR Purification Kit (Fermentas, Thermo Scientific, Waltham, MA, USA) according to the manufacturer's instructions. The concentration of DNA was measured using a NanoDrop 1000 Spectrophotometer (Nanodrop, Wilmington, DE, USA). The purified and quantified PCR products were sent to the Genome Quebec and McGill University Innovation Center for sequencing using Roche GS FLX Titanium technology.

For illumina sequencing, the quantified DNA samples were directly sent to Genome Quebec and McGill University Innovation Center for sequencing. The DNA samples were amplified using the modified primers: 926f modified (5'-AAACTYAAAKGAAT**W**G**R**CGG-3'), and 1392r modified (5'-ACGGGCGGTG**W**GTRC-3'), which were derived from previous publications (Ferris et al. 1996a; Engelbrektson et al. 2010).

##### iv. Dechlorination of HCH Isomers

Each HCH isomer can exist in two chair conformers which are in quick equilibrium at room temperature (Table S2). The β- and δ-HCH isomers have to undergo a ring-flip from the most stable chair conformer to an unstable conformer to obtain anti-parallel chorine atoms before the first dichloroelimination reaction. Although there is one axial chlorine on the δ-HCH molecule, the dehydrochlorination reaction is still impossible to initiate.

Abiotic dehydrochlorination reactions (eliminating HCl) can occur under alkaline conditions producing trichlorobenzene (TCB) isomers as the major end product (Cristol 1947; Hughes et al. 1953; Liu et al. 2003). The pH of the cultures in this study was kept ~6.9, and the final dechlorination products of HCH isomers were MCB and benzene (we never saw any TCB); therefore such consecutive abiotic dehydrochlorination reaction did not happen in our experiments. Rather, it is more likely that the four HCH isomers produced three tetrachlorocyclohexene (TeCCH) isomers as the intermediates.

The chlorine atoms in HCHs have two positions: axial or equatorial. When a carbon-carbon double bond forms (for example in TeCCH), the relative positions of the chlorines may change to achieve a structure with the lowest energy, however, tetrahedral carbons are still chiral and substituents are not free to rotate. As a result, the chlorine atoms in the TeCCH molecules are not in strictly axial/equatorial positions, but the two chlorines on the tetrahedral carbon are still in approximately axial/equatorial positions. We assumed that the chlorines in the “approximately” axial position are still more easily eliminated than those in similar equatorial positions.

Taking the δ-3,4,5,6-TeCCH generated by β-HCH as an example, the four chlorine atoms are in approximately coplanar equatorial positions, making the δ-TeCCH the most stable isomer. To further eliminate two chorines, the δ-TeCCH molecules have to ring-flip to obtain approximately anti axial position chlorine atoms to achieve possible maximum overlap orbitals when forming double bonds. However, for the γ-3,4,5,6-TeCCH produced by γ-HCH and δ-HCH, the Cl (6) is approximately axial to the ring while the other three chorines are on the approximately equatorial positions. From the perspective of minimizing energy, dehydrochlorination eliminating the axial chorine with the adjacent axial hydrogen to produce trichlorocyclohexadiene (TCCH,) is favored over dichloroelimination which requires a ring-flip to remove two chlorines.

**Table S1.** Selected physical and chemical properties of HCH isomers at 25°C [1]

| Property | Isomers | | | |
|---|---|---|---|---|
| | α | β | δ | γ |
| Water solubility (mg/L) | 1 | 0.1 | 8 | 7.3 |
| Solubility in 100 g ethanol [2] (g) | 1.8 | 1.1 | 24.4 | 6.4 |
| Vapor pressure (Pa) | 4.4E-02 | 4.0E-05 | 2.0E-03 | 3.7E-03 |
| Henry's law constant (Pa $m^3$/mol) | 0.872 | 0.116 | 0.0727 | 0.149 |
| $logK_{OW}$[3] | 3.8 | 3.8 | 4.14 | 3.7 |
| $logK_{oc}$[4] | 3.81 | 3.36 | 3.28 | 3.0 |

Note:

[1]All data were from (Mackay et al. 2006) unless otherwise noted.

[2]Data from (Clayton and Clayton 1981)

[3]$K_{OW}$: n-octanol/water partition coefficient

[4]$K_{oc}$: organic carbon-water partition coefficient.

β-HCH is far less soluble in water (<1 mg/L) than the other isomers, reflecting much more hydrophobic character and net dipole moment of zero. The vapor pressure and Henry's Law constant are also appreciably lower for the beta isomers as compared to the others. Collectively, these observations for β-HCH can be rationalized in terms of its compact, all-equatorial chlorine substituents and greater stability. The vapor pressure of α-HCH is approximately one order of magnitude higher than that of the other isomers. This elevated vapor pressure coupled with the large Henry's Law constant underscores the increased air partitioning behavior and substantial airborne transport risk. The $logK_{oc}$ values indicate the four isomers exhibit slight differences in sorption potential. As an example, the $logK_{oc}$ value reported for γ-HCH indicates that it should be the most soluble and least hydrophobic isomer, and consequently, exhibits the lowest sorption potential.

**Table S2.** Basic thermodynamic information on the two conformers of HCH isomers.

| Isomer | [1]Torsional strain in two conformers (kcal/mol) | | [2]$\Delta G^0$ (kcal/mol) | [3]Energy Barrier (kcal/mol) | [4]$k_{eq}$ | [5]Percent in stable conformer (%) |
|---|---|---|---|---|---|---|
| | Stable | Non-stable | | | | |
| α | 1.04 | 2.07 | 1.04 | 13.1 | 5.76 | 85.2% |
| β | 0 | 3.11 | 3.11 | 15.2 | 191 | 99.5% |
| δ | 0.52 | 2.59 | 2.07 | 14.2 | 33.2 | 97.1% |
| γ | 1.55 | 1.55 | 0 | 12.1 | 1 | 50% |

1. The two conformers of each HCH isomer are shown in Figure S1. The torsional strain of each chlorine atom on the axial position is 0.518 kcal/mol and the strain is additive (Bruice 2017).
2. The free energy differences between two chair conformers.
3. The energy barrier for interconversion between two chair conformers. The basic energy barrier for cyclohexane interconversion is 12.1 kcal/mol (Bruice 2017).
4. The equilibrium constant, $k_{eq}$, was calculated as $k_{eq} = \exp(-\frac{\Delta G}{RT})$.
5. The percentage of molecules in stable conformer at equilibrium was calculated as follows:

$$\%\ \text{of equatorial conformer} = \frac{k_{eq}}{k_{eq} + 1}$$

**Table S3** Feeding information (see excel file)

**Table S4.** Comparison of the ratios of Benzene to MCB with previous published values for the HCH isomers.

| [1]Isomer | [2]This study | Elango et al. 2011 | Doesburg et al. 2005 | Boyle et al. 1999 | Middeldorp et al. 1996 | Baker et al. 1985 |
|---|---|---|---|---|---|---|
| α (eeeeaa) | 0.55±0.09 (N=24) | | 0.3 | | | |
| β (eeeeee) | 0.77±0.15 (N=24) | | 0.7 | | 0.29 (85% recovery) | |
| γ (eeeaaa) | 0.13±0.02 (N=27) | 0.33 | 0.2 | 0.33 (60% recovery) | | 0.05 (40% recovery) |
| δ (eeeeea) | 0.06±0.02 (N=27) | | | | | |

1. The “e” and “a” in the brackets represent the equatorial and axial positions of chlorine atoms in the stable conformers, respectively.
2. Data are average (+/- stdev) of triplicate bottles for each isomer from Day 387 to Day 590.

**Table S5.** Number of sequences per sample during data processing

| Day | Sample name | Raw reads | # after joining | # after quality filtering | # after removal of chimera | | #of OTUs | |
|---|---|---|---|---|---|---|---|---|
| | | | | | All | No Archaea | All | No Archaea |
| 0 | Parent GT1 | | N/A | | 6068 | 4585 | 276 | 244 |
| 240 | α-1 | 83769 | 58702 | 56669 | 53805 | 19748 | 1785 | 978 |
| | α-2 | 69177 | 49026 | 46034 | 44520 | 15228 | 1061 | 596 |
| | α-3 | 71132 | 50329 | 47400 | 45193 | 14886 | 1343 | 743 |
| 595 | α-1 | 78553 | 52761 | 49310 | 45985 | 18898 | 1644 | 936 |
| | α-2 | 85909 | 59818 | 54917 | 51660 | 17162 | 1790 | 1058 |
| | α-3 | 79421 | 57075 | 53196 | 49965 | 11972 | 2162 | 1128 |
| 240 | β-1 | 70668 | 48323 | 45393 | 43314 | 16163 | 1355 | 777 |
| | β-2 | 72459 | 49265 | 46146 | 43877 | 17888 | 1474 | 850 |
| | β-3 | 74255 | 50252 | 47258 | 44949 | 17748 | 1390 | 823 |
| 595 | β-1 | 77840 | 54409 | 50540 | 49196 | 9981 | 1490 | 953 |
| | β-2 | 87101 | 61720 | 56805 | 55496 | 6901 | 1622 | 789 |
| | β-3 | 72839 | 51804 | 48655 | 46740 | 7043 | 1652 | 785 |
| 240 | γ-1 | 79821 | 55934 | 52373 | 49667 | 11524 | 2542 | 1486 |
| | γ-2 | 84555 | 58196 | 53218 | 50373 | 13269 | 2159 | 1393 |
| | γ-3 | 69713 | 48330 | 46610 | 44077 | 11099 | 2433 | 1444 |
| 595 | γ-1 | 88312 | 63042 | 59275 | 55987 | 17632 | 1620 | 937 |
| | γ-2 | 78060 | 55628 | 52107 | 49515 | 18903 | 1693 | 984 |
| | γ-3 | 90322 | 64017 | 59969 | 57455 | 9443 | 1406 | 851 |
| 240 | δ-1 | 68347 | 48294 | 45347 | 41814 | 12185 | 1795 | 828 |
| | δ-2 | 76916 | 54098 | 50640 | 47766 | 13311 | 1719 | 839 |
| | δ-3 | 70965 | 49573 | 46429 | 43135 | 10011 | 1623 | 736 |
| 595 | δ-1 | 82517 | 59309 | 55525 | 49904 | 14053 | 2071 | 890 |
| | δ-2 | 81560 | 58309 | 54581 | 49224 | 16467 | 1965 | 900 |
| | δ-3 | 91390 | 64024 | 59035 | 53955 | 13666 | 2069 | 843 |

**Table S6 (Excel).** qPCR reactions
**Table S7 (Excel).** Bacteria and Archaea identified in samples (normalized relative and absolute abundance calculations)
**Table S8 (Excel).** Yield calculations.

#### References


Amann, R.I., W. Ludwig, and K.-H. Schleifer. 1995. Phylogenetic identification and in situ detection of individual microbial cells without cultivation. *Microbiol. Rev.* 59 no. 1: 143-169.

Bruice, P.Y. 2017. *Organic Chemistry*. United States of America: Pearson Education, Inc.

Clayton, G.D., and F.E. Clayton. 1981. *Patty's industrial hygiene and toxicology*. 3rd ed: John Wiley & Sons.

Cristol, S.J. 1947. The Kinetics of the Alkaline Dehydrochlorination of the Benzene Hexachloride Isomers. The mechanism of second-order elimination reactions. *J. Am. Chem. Soc.* 69 no. 2: 338-342.

Engelbrektson, A., V. Kunin, K.C. Wrighton, N. Zvenigorodsky, F. Chen, H. Ochman, and P. Hugenholtz. 2010. Experimental factors affecting PCR-based estimates of microbial species richness and evenness. *ISME J.* 4 no. 5: 642-647.

Ferris, M., G. Muyzer, and D. Ward. 1996a. Denaturing gradient gel electrophoresis profiles of 16S rRNA-defined populations inhabiting a hot spring microbial mat community. *Appl. Environ. Microbiol.* 62 no. 2: 340-346.

Ferris, M.J., G. Muyzer, and D.M. Ward. 1996b. Denaturing gradient gel electrophoresis profiles of 16S rRNA-defined populations inhabiting a hot spring microbial mat community. *Applied & Environmental Microbiology* 62 no. 2: 340.

Hanshew, A.S., C.J. Mason, K.F. Raffa, and C.R. Currie. 2013. Minimization of chloroplast contamination in 16S rRNA gene pyrosequencing of insect herbivore bacterial communities. *J. Microbiol. Methods* 95 no. 2: 149-155.

Hughes, E.D., C.K. Ingold, and R. Pasternak. 1953. Mechanism of elimination reactions. Part XVIII. Kinetics and steric course of elimination from isomeric benzene hexachlorides. *Journal of the Chemical Society*: 3832-3839.

Kocur, C.M., L. Lomheim, O. Molenda, K.P. Weber, L.M. Austrins, B.E. Sleep, H.K. Boparai, E.A. Edwards, and D.M. O'Carroll. 2016. Long-term field study of microbial community and dechlorinating activity following carboxymethyl cellulose-stabilized nanoscale zero-valent iron injection. *Environ. Sci. Technol.* 50 no. 14: 7658-7670.

Liu, X., P.a. Peng, J. Fu, and W. Huang. 2003. Effects of FeS on the Transformation Kinetics of γ-Hexachlorocyclohexane. *Environmental Science & Technology* 37 no. 9: 1822-1828.

Mackay, D., W.Y. Shiu, K.-C. Ma, and S.C. Lee. 2006. *Physical-chemical properties and environmental fate for organic chemicals.* 2nd ed: CRC Press.
